# No evidence for short-term evolutionary response to a warming environment in Drosophila

**DOI:** 10.1101/2021.02.19.432014

**Authors:** Marta A. Santos, Ana Carromeu-Santos, Ana S. Quina, Mauro Santos, Margarida Matos, Pedro Simões

## Abstract

Adaptive evolution is key in mediating responses to climate change. Such evolution will expectedly lead to changes in the populations’ thermal reaction norm and improve their ability to cope with stressful conditions. Conversely, constraints of different nature might limit the adaptive response. Here, we test these expectations by performing a real-time evolution experiment in historically differentiated *Drosophila subobscura* populations. We address the phenotypic changes of flies evolving for nine generations in a daily fluctuating environment with average constant temperature, or a warming environment with increasing average and amplitude temperature across generations. Our results showed that (1) evolution under a global warming scenario has not led, so far, to a noticeable change in the thermal response; (2) historical background appears to be affecting the responses of populations under the warming environment, particularly at higher temperatures; (3) thermal reaction norms are trait-dependent: while lifelong exposure to low temperature decreases fecundity and productivity but not viability, high temperature causes negative transgenerational effects on productivity and viability, even though fecundity remains high. These findings raise concerns about the short-term efficiency of adaptive responses to the current changing climate.

## 1. Introduction

One catastrophic consequence of climate change is the global decimation of biodiversity, which impacts on species abundance and distribution and damages the ecosystem functioning (Somero 2012; Pecl et al. 2017; IPCC 2018). Ectotherms are a huge fraction of biodiversity, have an essential role in ecosystems, and are particularly vulnerable to global warming due to the profound effects of temperature on their biology, biochemistry, and physiology (e.g. Dillon et al. 2010; Hoffmann and Sgrò 2018). In species with low dispersal ability or when migration to climatically appropriate habitats is not possible, adaptation to the changing environmental conditions is crucial for survival (Hoffmann and Sgró 2011; Huey et al. 2012). Recent evidence suggests that adaptation to higher temperatures might be highly limited by physiological constraints and/or possible lack of genetic variation (Kellermann et al. 2009, 2012; Araújo et al. 2013; Kristensen et al. 2015). It is, thus, crucial to understand the potential of species to adaptively respond to global warming, and the underlying molecular, physiological, and phenotypic changes (Franks and Hoffmann 2012; Urban et al. 2016; Bell 2017). This will shed light on how evolution shapes the populations’ fitness and fate, increasing our ability to preserve biodiversity and forecast species distributions (Urban et al. 2016; Bay et al. 2018).

Thermal environments are often quite dynamic in nature, with both circadian and seasonal variation, in addition to increasing mean temperatures and extreme thermal events as a consequence of global warming (IPCC 2018). There is increasing awareness of the impact of thermal variation on organism fitness (Bozinovic et al. 2011; Vasseur et al. 2014; Colinet et al. 2015; Cavieres et al. 2018), a key aspect that needs to be taken into account in experimental studies addressing thermal adaptation, in general, and evolutionary potential to respond to climate change, in particular (Hallsson and Björklund 2012; Schou et al. 2014; Manenti et al. 2016; Sørensen et al. 2020). The amount and rate of evolutionary change in response to climate change is dependent on diverse aspects such as the levels of additive genetic variation, trade-offs derived from different thermal physiology (within and across species), population size and structure, inbreeding, and the rate of environmental change (Chevin et al. 2010; Hoffmann and Sgró 2011; Bell 2017; Kristensen et al. 2018; Trubenová et al. 2019). One important body of theory that allows envisioning how adaptation to thermally varying environments proceeds stems from the generalist *vs*. specialist trade-off. Specialists are predicted to evolve in the more stable thermal environments and generalists in the more dynamic, variable environments (Angilletta et al. 2003; Angilletta 2009). These predictions assume a trade-off between maximal performance and performance breath, resulting from antagonistic pleiotropy associated with constrains to the structure and function of enzymes at different temperatures (Huey and Kingsolver 1989; Angilletta et al. 2003). Important empirical work has been done on evolution under constant *vs*. variable thermal environments, in controlled laboratory experiments. Overall, the expectations of a specialist *vs*. generalist trade-off have not been supported (Ketola et al. 2013; Berger et al. 2014; Condon et al. 2014; Manenti et al. 2016; but see Le Vinh Thuy et al. 2016). However, there is some consistent evidence for the evolution of generalists under fluctuating environments (Ketola et al. 2013; Condon et al. 2014). The occurrence of indirect selection, the range of tested temperatures (Condon et al. 2015), and the estimation of performance based on constant rather than fluctuating temperatures (Ketola and Saarinen 2015) might explain, at least in part, why evidence for a specialist *vs*. generalist trade-off is lacking. Alternatively, other mechanisms might govern the evolution of thermal performance, namely, through allocation and acquisition trade-offs, in which one given phenotype might outperform other across a wide range of temperatures (Angilletta et al. 2003).

The study of real-time evolution of populations subjected to different realistic ecological scenarios can also provide invaluable insight on the evolution and adaptive potential to respond to environmental fluctuations and climate change (Kawecki et al. 2012; Bailey and Bataillon 2016; Kellermann and van Heerwaarden 2019). By means of experimental evolution, populations are studied across several generations in very well defined and reproducible conditions that are often achieved only in the laboratory (*e*.*g*., Kawecki et al. 2012; Magalhães and Matos 2012). This powerful approach can provide direct evidence for adaptation to diverse thermal environments in different key adult and juvenile traits, allow the estimation of adaptive change rates, and elucidate the link between phenotypic and genetic variation (Hoffmann and Sgró 2011; Porcelli et al. 2015). Experimental evolution has been a tool of choice to address several mechanisms and dynamics of thermal evolution in the last fifteen years (*e*.*g*., Santos et al. 2005, 2006; Hallsson and Björklund 2012; Rogell et al. 2014; Schou et al. 2014; Kellermann et al. 2015; Tobler et al. 2015; Manenti et al. 2016; Kinzner et al. 2019). Some of these studies have focused on evolution of ectotherms under increasing temperatures, as expected in a global warming scenario, although with varying rates of environmental change (Hallsson and Björklund 2012; Rogell et al. 2014; Schou et al. 2014; Kinzner et al. 2019). Most of them provided evidence of limited potential for evolutionary responses (Hallsson and Björklund 2012; Schou et al. 2014; Kinzner et al. 2019)). However, no empirical studies have yet addressed, to our knowledge, the impact on the evolutionary response of increasing both thermal mean and amplitude, both key aspects of global warming (IPCC 2018).

*Drosophila* is an excellent model organism to study thermal adaptation in ectotherms and has been widely used in experimental evolution studies. Disparities among studies have been obtained and highlight that thermal adaptive responses are complex and caused by multiple factors, particularly, the specific thermal environments and populations under study; also, different methodologies to estimate stress response contribute to the different conclusions (Hoffmann and Sgró 2011; Kellermann and van Heerwaarden 2019; Kristensen et al. 2020). *Drosophila subobscura*, a native Palearctic species, is a well-known case study of thermal adaptation. It shows genetic variation associated with local adaptation, with evidence of latitudinal clinal variation for chromosomal inversion frequencies in Europe and, more recently, in other two continents after colonization of South and North America (Rezende et al. 2010). These polymorphisms have also been shifting worldwide, associated with global warming (Balanyá et al. 2006) and responding to heat waves (Rodríguez-Trelles et al. 2013). Previous selection experiments using different temperatures showed that evolutionary responses were not always as expected from clinal patterns (Santos et al. 2005). This species presents relevant thermal plasticity, with developmental temperature playing a decisive role in adult reproductive performance (Simões et al. 2020; Santos et al. 2021). The range of development temperatures suitable for *D. subobscura* is 6 – 26 °C (Moreteau et al. 1997; David et al. 2005; Schou et al. 2017) with optimal viability between 16 °C and 20°C (Schou et al. 2017), which is in agreement with the preferred body temperature (Rego et al. 2010; Castañeda et al. 2013). This species displays clear plastic responses to new thermal challenges (Fragata et al. 2016; Simões et al. 2020; Santos et al. 2021), with some evidence for historical differences between populations, particularly at colder temperatures (Simões et al. 2020).

Here, we use experimental evolution to analyze the evolutionary response of historically differentiated *D. subobscura* populations, derived from extreme latitudes of the European cline (Simões et al. 2017, 2020), after nine generations under different thermal selective regimes that comprise differences in mean temperature and/or thermal amplitude. These regimes include (1) a constant thermal environment, corresponding to the thermal conditions of the long-established populations (controls), (2) a circadian thermal fluctuating environment (cooler nights, warmer days), and (3) a global warming-like environment, with increases in thermal mean and amplitude across generations (progressively lower and higher thermal extremes across generations) – Figures 1a and 1b. Adult reproductive performance of populations evolving under each of these environments was tested in different combinations of juvenile and adult temperatures. This experimental setup allowed us to address the effects of thermal fluctuations *per se*, and of a warming environment on the populations’ evolutionary response. We have three general expectations for this response. First, populations evolving under more heterogeneous (fluctuating) thermal conditions are expected to evolve an overall better performance across different thermal environments (*i*.*e*., jack-of-all-temperatures, Huey and Kingsolver 1989). Second, populations evolving in warming conditions will, most likely, perform better in more stressful thermal environments, as these populations experience such temperatures during their life cycle. Third, assuming there are costs of adaptation to more extreme temperatures (Huey and Kingsolver 1989; Angilletta et al. 2003), populations under fluctuating and (even more so) warming conditions will perform worse in intermediate thermal conditions than the control populations. Alternatively, lack of genetic variation or trade-offs between traits might limit the thermal evolutionary response. We therefore aimed to respond to the following questions: (i) Does evolution under warming or fluctuating environments change the population’s thermal reaction norm? (ii) Does lifelong (or adulthood only) exposure to more extreme temperatures lead to a decline in performance? If so, does evolution in a warming environment change performance under these stressful conditions? (iii) Can we find evidence of geographical differences in the populations’ thermal evolutionary response?

**Figure 1a.**
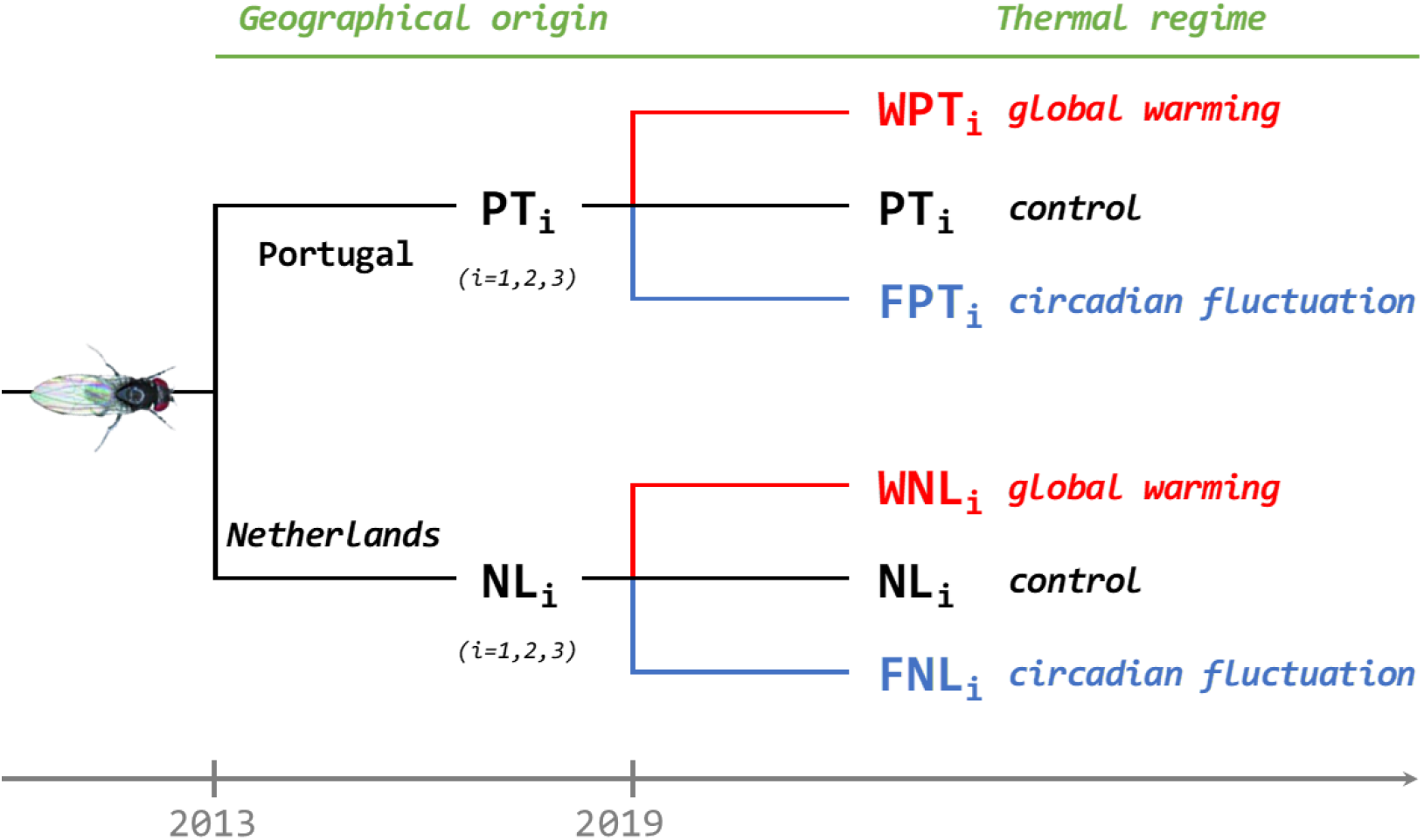
Derivation of the experimental lines from natural Drosophila subobscura populations.

**Figure 1b.**
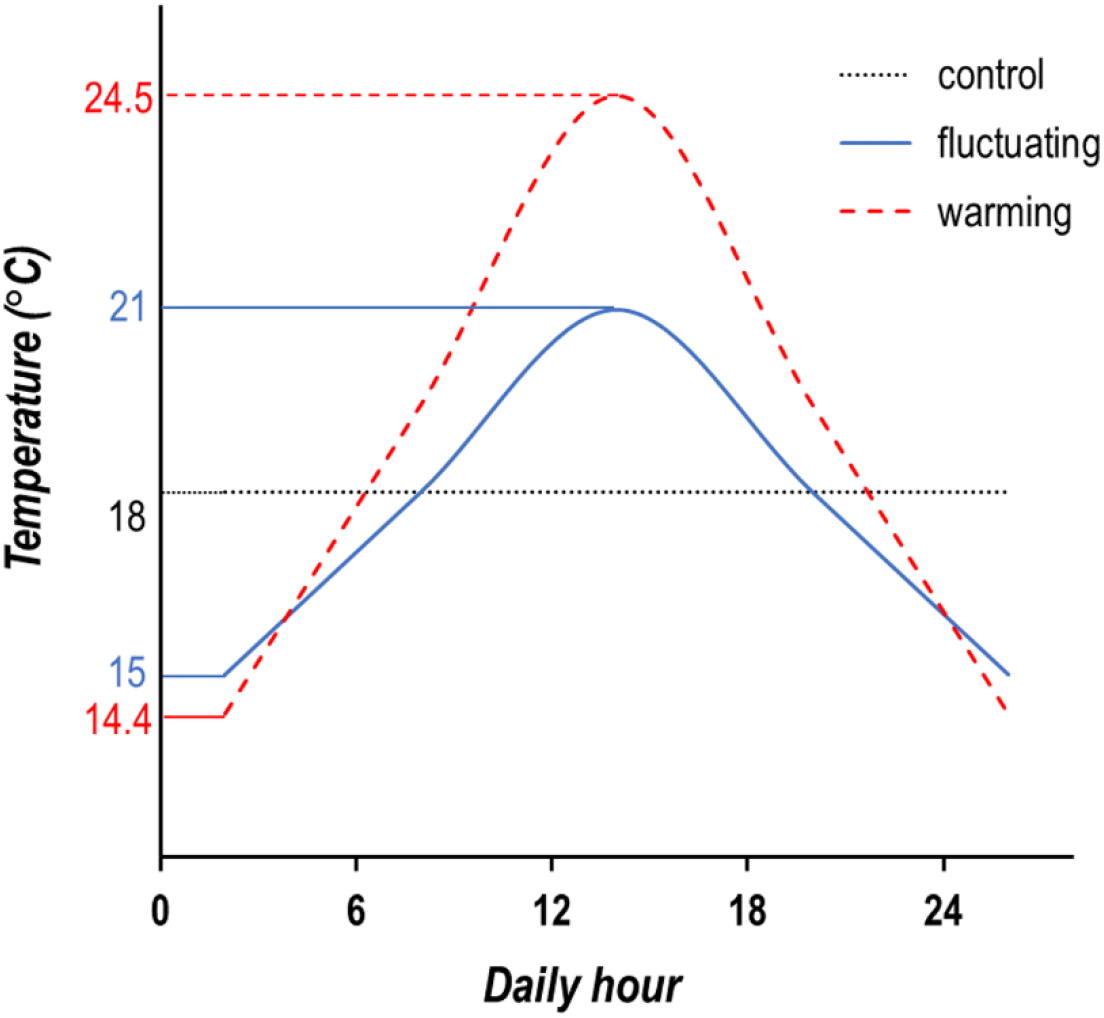
Daily temperature profiles of the experimental regimes after nine generations of thermal selection.

## 2. Material and Methods

### 2.1. Laboratory Populations and Thermal Selection Regimes

In late August/early September 2013, two natural populations of *Drosophila subobscura* were sampled. The collections were done in Adraga, Portugal (lat. 38°48′ N) and Groningen, The Netherlands (lat. 53°13′ N); two contrasting latitudes of the European cline (ranging from Scandinavia, ~60°N, to Northern Africa, ~30°N; Prevosti et al. 1988) that experience distinct environmental temperatures. From these samples two sets of laboratory populations were established: PT, from Adraga, and NL, from Groningen – see details in (Simões et al. 2017). Each population was three-fold replicated in the lab, originating PT_1-3_ and NL_1-3_ populations. Maintenance involved discrete generations with a synchronous 28-day cycle and discrete generations, 12L:12D photoperiod, constant temperature of 18°C, controlled densities in both adults (50 adults per vial) and eggs (70 eggs per vial) in ~30 mm3 glass vials, and reproduction for the next generation around peak fecundity (seven to ten days old imagoes). Census size ranged between 500 and 1000 individuals (see Simões et al. 2017). In January 2019, after the PT and NL populations had undergone 70 generations of lab evolution, two new thermal selection regimes were derived (Figure 1a): *circadian fluctuation* (F, originating FNL_1-3_ and FPT_1-3_) and *global warming* (W, originating WNL_1-3_ and WPT_1-3_).

The F regime is under a daily temperature fluctuation between 15°C and 21°C, with a mean daily temperature of 18°C and constant across generations. The W regime has a daily fluctuation similar to the F regime, but has a 0.2°C increase in daily mean and 0.5°C increase in daily amplitude (difference between the highest and lowest temperature) every generation. After nine generations of thermal selection, when the experiment was carried out, the Warming populations have been subjected to temperatures ranging from 14.4°C to 24.5°C, with a daily average of 19.4°C. (Figure 1b). By generation seven, the temperature increase has led to a 24h reduction in the life-cycle length, which became 27 days. The PT and NL populations, kept at constant 18°C, are the experimental controls, representing the ancestral state of the other two thermal regimes and accounting for environmental effects across generations. All experimental populations were, otherwise, maintained under the experimental protocol referred to above for the NL and PT populations. Census sizes were generally high for all thermal selection regimes (see Figure S1; average census for W= 935.6; F= 976.5 and C= 1006.1).

### 2.2. Experimental assays

To study the effect of different thermal environments on the performance of the experimentally evolved populations, we analyzed the performance of our eighteen experimental populations subjected to five thermal treatments (Figure 2). In three of these treatments, flies were exposed to the same developmental and adult temperatures, either a colder temperature (14°C), intermediate (18°C), or warmer temperature (24°C) – 14-14, 18-18, and 24-24 treatments, respectively. Because we previously had detected reduced performance under lower adult temperatures and higher developmental temperatures (Simões et al. 2020; Santos et al. 2021), two additional and potentially stressful thermal treatments were tested: 18-14 (lower temperatures in the adult stage) and 24-18 (higher temperatures during the developmental stage). To minimize maternal effects, the experiment was preceded by one generation of common garden in control conditions (18°C).

**Figure 2.**
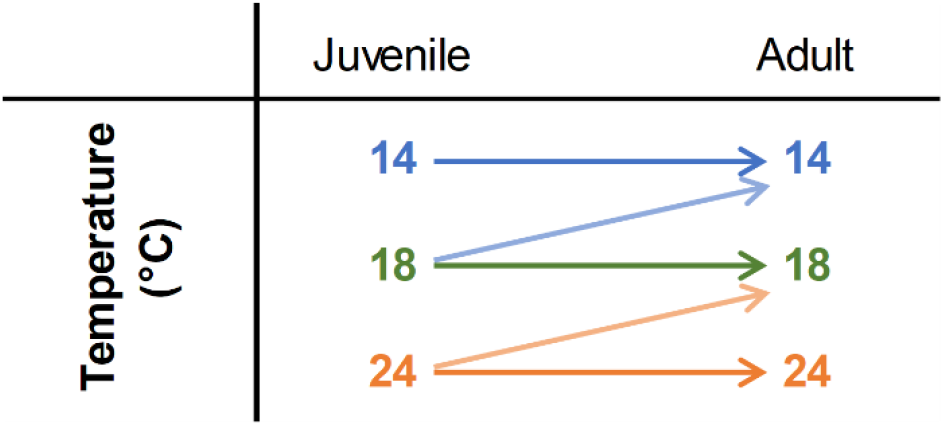
Experimental design: combinations of three developmental and three adulthood test temperatures.

Sixteen recently emerged mating pairs (virgin males and females) per population and treatment were formed, with a total of 1440 pairs (16 pairs × 18 populations × 5 temperature treatments). Flies were transferred to fresh medium every other day, the vials were daily checked for the presence of eggs, and the eggs laid by each female were counted between days six and eight since emergence. Four life-history traits were studied. (1) *age of first reproduction* (number of days since emergence until the first egg laying), which addresses the rate of sexual maturity; (2) *fecundity* (total number of eggs laid between days six and eight), which refers to a period that is close to the age of egg collection for the following generation (seven to ten-day-old imagoes), where selective pressures are likely higher; (3) *productivity* (number of emerged flies from the eggs laid on day eight), which shows the ability of a female to produce viable descent, and (4) *juvenile viability* (ratio between productivity and fecundity at day eight), which conveys to the efficiency of reproduction. For reliability purposes, only vials with at least five laid eggs were considered for juvenile viability.

### 2.3. Statistical Methods

Raw data used in the analyses is the mean value for each replicate population and temperature treatment (*e*.*g*., the mean of PT_1_ for the 14-14 treatment is one of the three data values for PT in this treatment). Data was analyzed by linear mixed effects models fitted with REML (restricted maximum likelihood). p-values for differences between temperature treatments, thermal regimes, populations as well as their interactions were obtained through analyses of variance (Type III Wald F tests, Kenward-Roger degrees of freedom). Two general models were applied (for simplicity we do not present interactions with random factors):

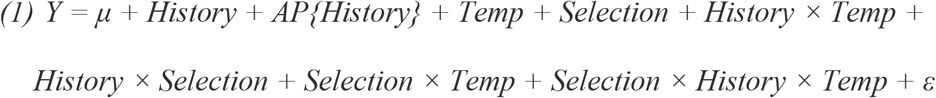

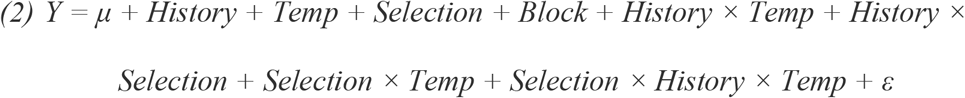

where Y is the studied trait (age of first reproduction, fecundity, productivity, or viability), History is the fixed factor corresponding to distinct geographical origin (with categories PT and NL), Selection is the fixed factor representing the Thermal Selection Regimes (with three levels: Control, Warming, and Fluctuating), and Temp is the fixed factor corresponding to the five different temperature treatments. In model (1), AP{History} is the random effect consisting of the ancestral replicate population (*i*.*e*., PT1-3; NL1-3) nested in History, from which each of the replicate populations of the three thermal selection regimes were generated (*e*.*g*., Ancestral PT1 originating Control PT1, Fluctuating PT1 and Warming PT1). In model (2) Block was defined as random effect, corresponding to the set of same-numbered replicate populations from all different thermal regimes that were assayed synchronously in the same experimental rack in a randomized manner. Models with and without interactions with random factors were assessed using AIC and the best model for each trait was chosen: model (1) for age of first reproduction and fecundity including interactions with AP; model (2) for productivity, without defining interactions with the random block. For the viability data, models (1) and (2) were tested including fecundity of day eight (F8) as covariate to account for the variation in fecundity across thermal treatments. The model without interactions with AP (random factor) and defining F8 as covariate presented the lowest AIC and was, therefore, chosen.

This body of data allowed to address how differences in the experimental populations associated with thermal regime (Selection) and geographical origin (History) impacted on their thermal performance. First, comparisons between thermal regimes were performed for the lifelong thermal treatments (14-14 *vs*. 18-18 *vs*. 24-24), allowing to analyze the populations’ thermal reaction norms. Specific paired combinations of thermal regimes were used to test for (1) the effect of evolution under thermal fluctuations (F *vs*. C); and (2) the effect of evolution under global warming (W *vs*. C).

Second, specific effects of different temperature combinations were assessed to analyze how thermal evolution impacted on the flies’ performance under stressful environments. Two models were then applied to compare the Warming and the Control regimes: (1) under colder conditions (treatments 14-14, 18-14, and 18-18); and (2) under warmer conditions (treatments 24-24, 24-18, and 18-18). The factor History and its interaction with temperature were also included in these models to test for differences between populations with distinct biogeographical origin (PT vs. NL). Tukey post-hoc tests were used to compare performance in the three different thermal treatments, whenever the temperature treatment factor was significant, allowing to identify which thermal treatments showed reduced performance. We further tested for differences in performance due to selection (W *vs*. C) in the traits that presented a significant temperature effect, either for colder or warmer temperatures. In the latter analyses False Discovery Rate (FDR, Benjamini and Yekutieli 2001)) was applied to correct for multiple testing.

The homoscedasticity and normality assumptions for analysis of variance were checked and met in our dataset. Arcsine transformation was applied to the viability data to meet normality assumptions. All statistical analyses were performed in R v4.0.0, with *lme4* (Bates et al. 2015), *car* (Fox and Weisberg 2019), *lawstat* (Hui et al. 2008), *emmeans* and *ggplot2* (Wickham 2016) packages.

## 3. Results

### 3.1. Evolutionary Response under dynamic thermal environments

We tested the evolutionary response of populations evolving under the following dynamic thermal environments: (1) circadian fluctuating and (2) global warming, by comparing the performance of each thermal regime with the controls (Figures 3 and 4, respectively). These analyses included the life-long temperature test environments (14-14, 18-18, and 24-24). Both fluctuating *vs*. control and warming *vs*. control analyses showed differences between temperature treatments for all traits (significant temperature factor; 0.001<p<0.01 for fecundity and p<0.001 for all other traits in both comparisons, see Tables S1 and S2). However the patterns differed between them. For the fecundity characters (*i*.*e*., age of first reproduction and fecundity), a reduction of performance was only observed at lower temperatures (14-14 treatment), with a similar performance at control (18-18) and warmer (24-24) conditions. As for productivity, both colder and warmer temperatures showed a lower performance relative to control conditions. Viability showed yet a different pattern, with lower performance occurring only at the warmer temperature (see Figures 3 and 4).

**Figure 3.**
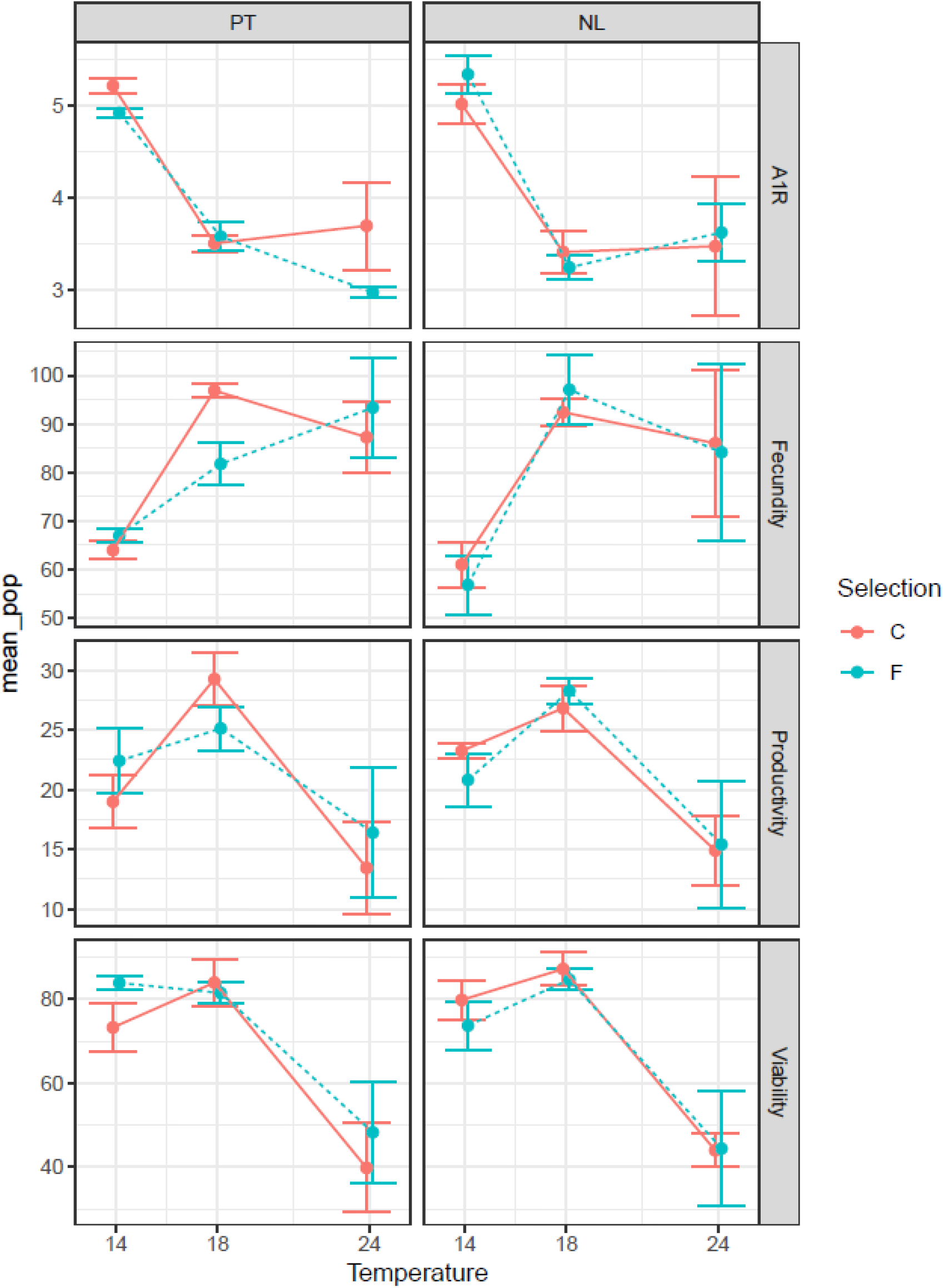
Thermal reaction norms of the Fluctuating (F) and Control (C) thermal selection regimes. A1R – Age of first reproduction. Values shown are the average and standard error for each thermal regime (using average values of each replicate population as raw data).

**Figure 4.**
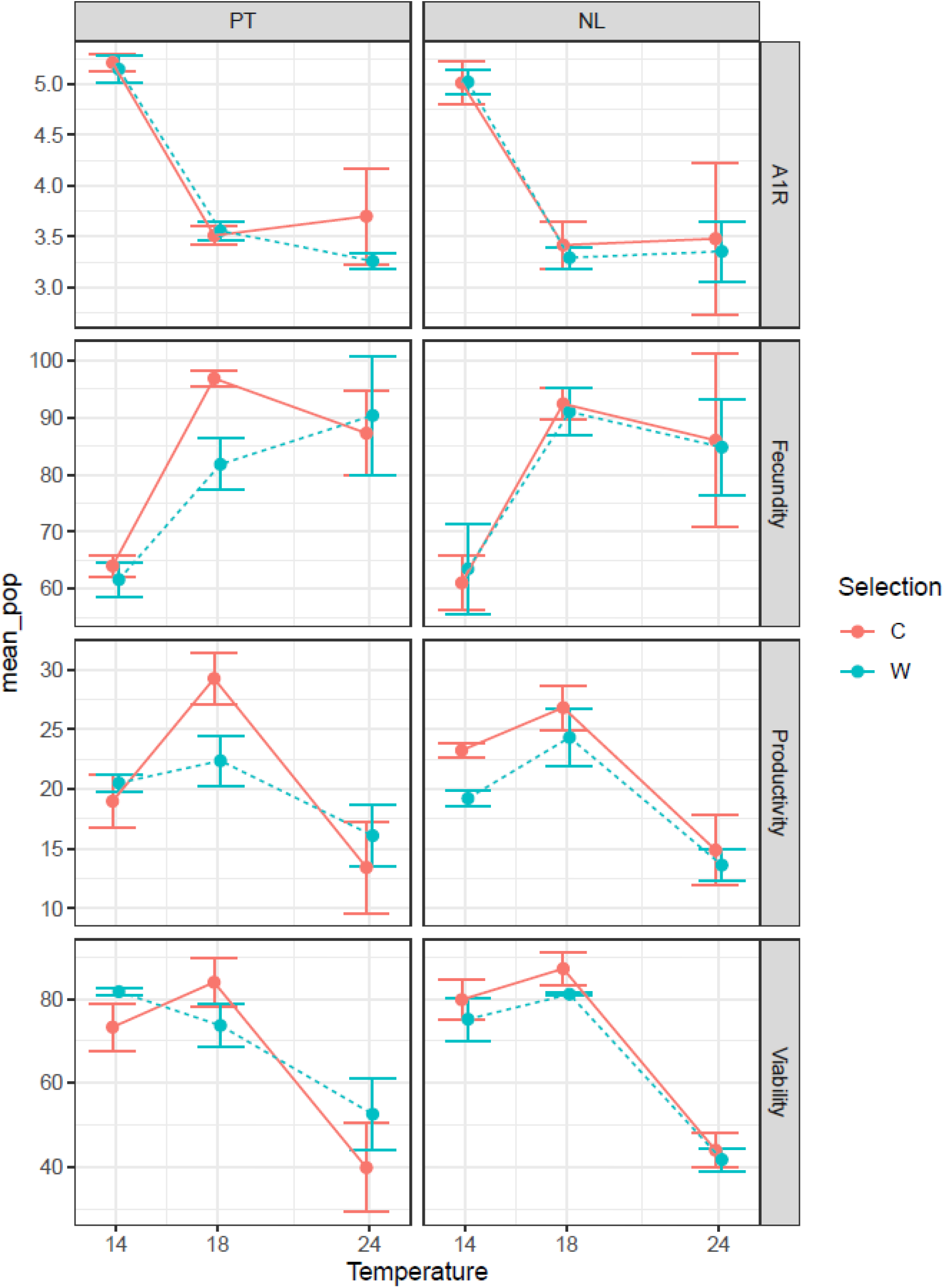
Thermal reaction norms of the Warming (F) and Control (C) thermal selection regimes. A1R – Age of first reproduction. Values shown are the average and standard error for each thermal regime (using average values of each replicate population as raw data).

The reaction norms of populations from the fluctuating or warming regimes were not significantly different from those of the controls (selection*temperature interaction, see Tables S1 and S2; Figures 3 and 4). Also, the reaction norms did not differ between populations with distinct histories (Portugal *vs*. Netherlands) – history*temperature interaction. Finally, the overall effects of history or selection were also not significant for any trait (see Tables S1 and S2).

### 3.2. Evolutionary response under stressful conditions

We aim to test whether populations adapting to a warming environment present an increased performance under stressful thermal environments. With this goal in mind, we assessed whether different juvenile-adult temperature treatments showed indications of cold stress (14-14, 18-14, and 18-18) or heat stress (24-18, 24-24, and 18-18), by comparing data from warming and control thermal selection regimes. Significant differences in performance between temperature treatments were obtained in both models – testing for either cold or heat stress – although this response varied across traits (see Table 1, Figures 4, S2 and text below). The exception was viability that did not present significant effect of lower temperatures.

**Table 1.**
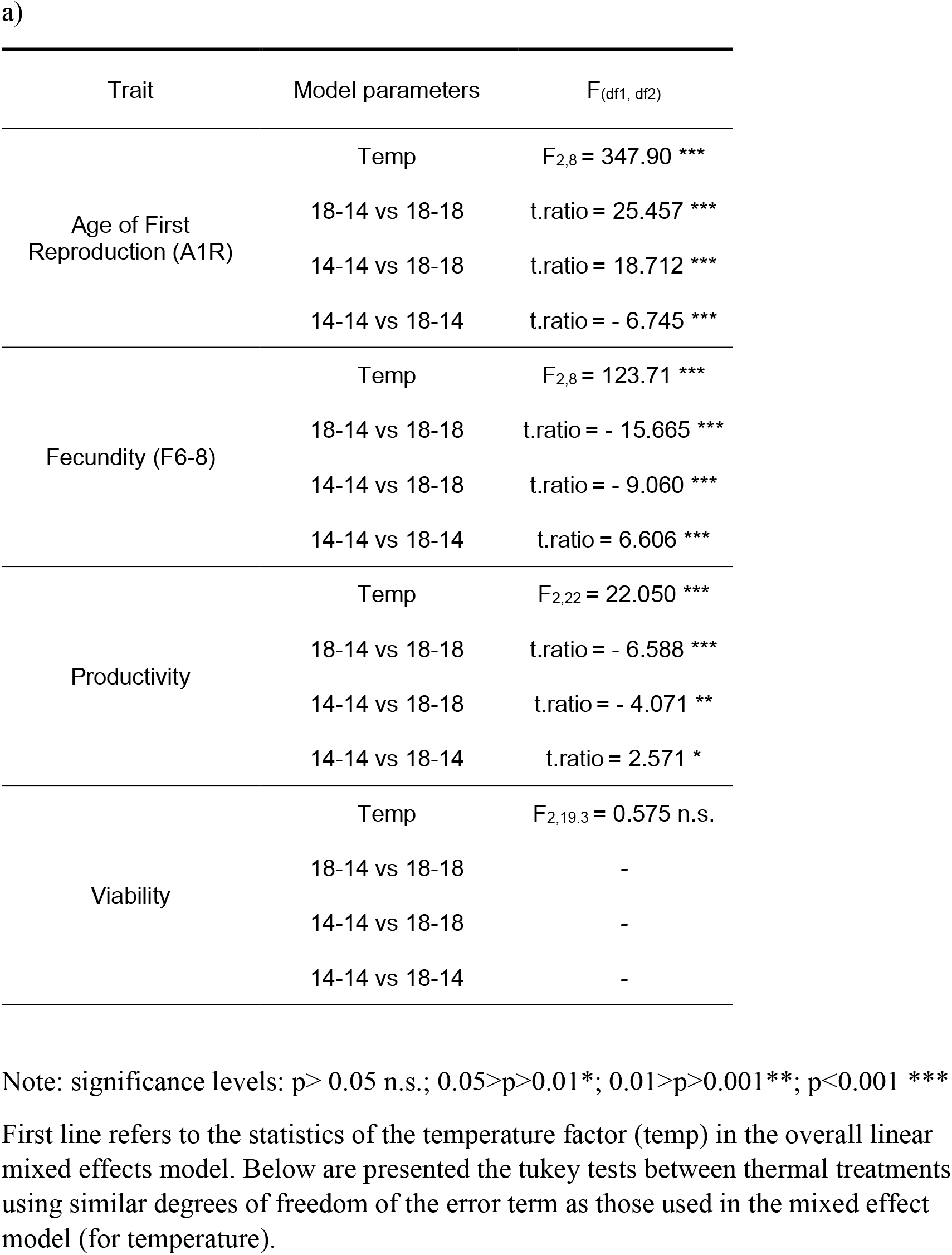
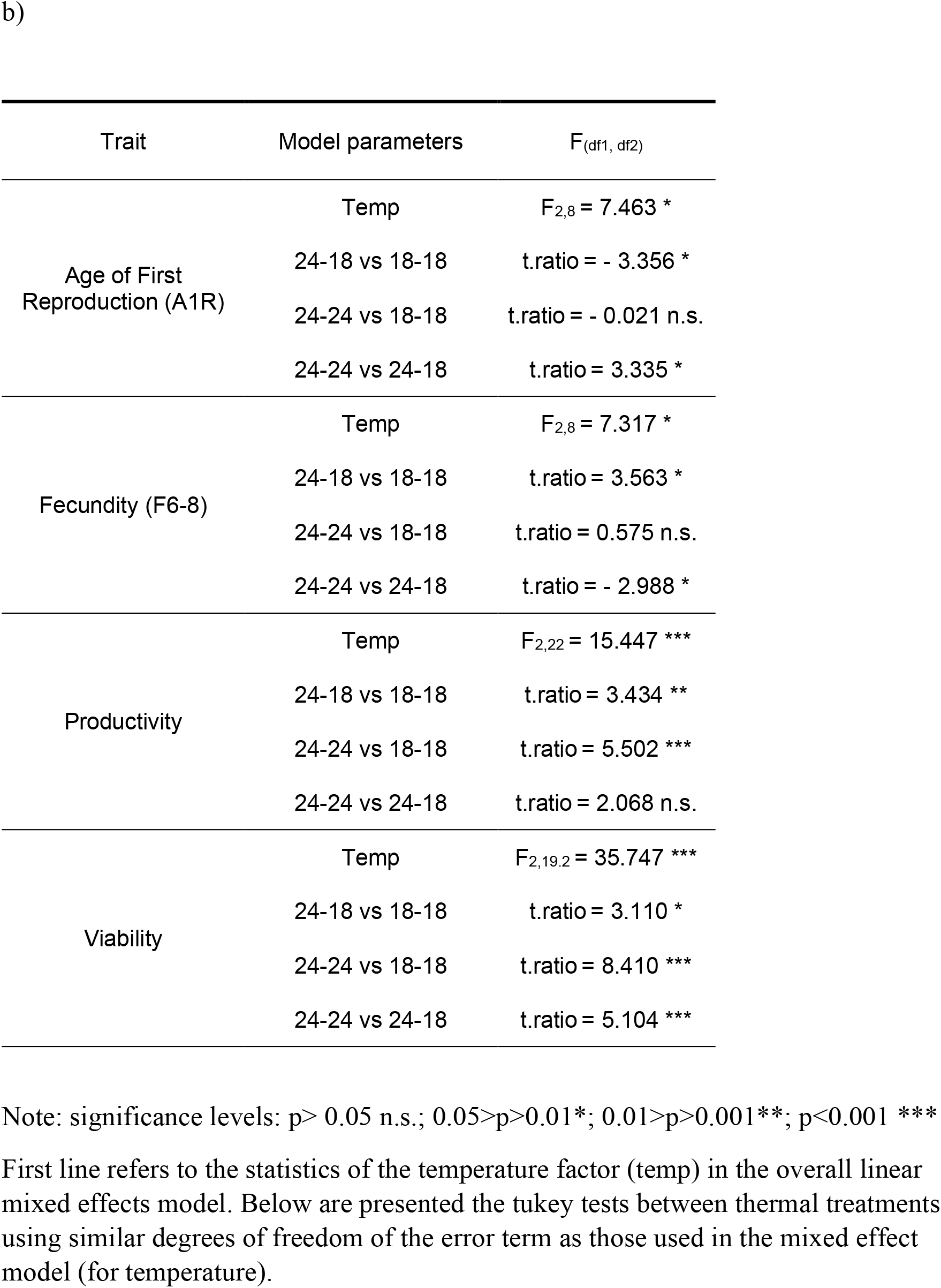
Differences in thermal performance between warming and control populations for colder (a) or warmer (b) temperature treatments.

When testing for heat stress, fecundity characters (*i*.*e*., age of first reproduction and fecundity) showed a reduction of performance when populations developed at 24°C were exposed to 18°C as adults (24-18 treatment) relative to those under the 18-18 and 24-24 treatments (see Table 1, Figure S2). However, no differences were found in performance between the 24-24 treatment and the control (18-18) conditions (Table 1). As for productivity and viability, both 24-24 and 24-18 show lower performance relative to control conditions. Interestingly, no significant differences in productivity were found between these warmer treatments, while viability was significantly lower in the 24-24 treatment relative to the 24-18 conditions – see Table 1 and Figure S2.

As for the cold stress analysis, significant reduction in performance was found in both colder treatments (14-14 and 18-14) relative to control conditions for age of first reproduction, fecundity, and productivity – see Table 1. For these same traits, performance of flies in the 14-14 conditions was always significantly higher than those of the 18-14 treatment. Viability showed no significant differences between these thermal treatments.

The linear mixed effects models applied to the pairwise combinations of temperature treatments showed no significant effects of selection or selection*temperature, in either cold or warm scenarios (see Table S3), which suggests that both warming and control populations do not differ in their response under the tested conditions. Interestingly, for the warming regime, we observed that southern populations (WPT_1-3_) presented consistently higher average values than their northern counterparts (WNL_1-3_) for all analyzed traits when tested in the warmer conditions (24-18 and 24-24; see Figures 4 and S2). However, no significant effects of biogeographical history or its interactions with temperature and selection were found for either heat or cold stress for any trait (Table S3).

## 4. Discussion

Recent empirical data casts doubt on the capacity of ectotherms, including *Drosophila*, to respond to climate change, with stress resistance traits being evolutionarily constrained (reviewed in (Kristensen et al. 2018; Kellermann and van Heerwaarden 2019). This is very worrisome, given the fast-adaptive demand caused by the rapid and globally changing environment. On the bright side, some thermal selection experiments have demonstrated that heat tolerance can evolve, when directly selected upon (Bubliy and Loeschcke 2005; Hangartner and Hoffmann 2016), although the ecologically relevance of such selection protocols might be questioned. Using the terminology of Diamond and Martin (2020), the question stands: can thermal adaptation be regarded as the silver bullet to fight global warming? According to our experimental data, the quick answer is no, not in the short run at least.

### Evolution in dynamic thermal environments did not substantially change life-history reaction norms

Nine generations of dynamic thermal evolution did not alter the thermal reaction norms of our experimental populations, nor did adaptation to a global warming environment improve their ability to cope with more stressful conditions. This lack of a clear adaptive response has been found in other studies addressing short-term evolution (< 20 generations) of ectotherms in increasingly warmer fluctuating environments, although with constant thermal amplitudes (Hallsson and Björklund 2012; Schou et al. 2014; Kinzner et al. 2019). In addition, evolution of *Drosophila simulans* populations in fluctuating environments with a constant average led to changes in the elevation of thermal reaction norms (increased mean performance across environments) but not in their shape (Manenti et al. 2015).

Several reasons could account for the deviation from our initial expectations. First, the number of elapsed generations might have been too small for evolutionary response to be detected. Because life-history traits and resistances to environmental stress are, in general, polygenic traits with low-to-intermediate heritabilities (Mousseau and Roff 1987; Diamond 2017; Castañeda et al. 2019), the evolutionary rates of adaptation are expected to be slow (Hoffmann et al. 2017). This might be the case even with a high rate of environmental change (Chevin et al. 2010; Hoffmann and Sgró 2011; Kristensen et al. 2018), as occurs in the warming environment we imposed – an increase of 0.2°C per generation, with a final daily peak temperature almost 7°C above control conditions.

Second, environmental stress can lead to population bottlenecks, which may dramatically decrease effective population size with the consequent loss of genetic variation and evolutionary response impairment (Hoffmann et al. 2017). However, demographic data from the experimental lines did not show any relevant census sizes drops (Figure S1). Furthermore, the control populations (in which this experiment was based) were found to have abundant genomic variation at least during the first 26 generations after lab founding (data not shown). This is corroborated by a previous study from our team, using lab populations collected from the same geographical locations, where only a very modest decline in genome diversity after 50 generations of lab evolution was found (Seabra et al. 2018). In addition, inversion polymorphism is known to have an adaptive response to thermal changes in this species (*e*.*g*., Rezende et al. 2010). A total of thirteen different inversions were still segregating in these populations after 68 generations in the laboratory (unpublished data). This level of polymorphism was already found by Santos et al. (2016) in other laboratory populations. Other *Drosophila* experiments involving increasing temperatures did not find an overall decline in genetic diversity (Schou et al. 2014; Kinzner et al. 2019), although we cannot exclude that low additive genetic variation for the analyzed phenotypic traits might be a factor in both those studies and ours.

Third, the idiosyncrasies of the dynamic thermal regimes, such as the simultaneous selection for lower and higher thermal extremes, may have slowed down the evolutionary response due to the existence of unfavorable genetic correlations and trade-offs. Though Manenti et al. (2016), in a study of evolution of *D. simulans* in fluctuating environments, found significant negative correlations between life-history and stress resistance traits, such trade-off between measures of cold and heat tolerance was not observed. Nevertheless, in a fluctuating environment, the time spent in each environment is necessarily shorter than in constant conditions, which reduces the strength of selection for each given environment (Kristensen et al. 2018). Finally, there might be some detachment between the dynamic selection environment and the conditions at which the performance tests were carried out (Ketola and Saarinen 2015). Further planned experiments will include, alongside with the thermal reaction norm approach, testing environments that match more precisely the conditions in which populations evolved. Interestingly, all southern populations of the warming regime (WPT) performed consistently better at warmer temperatures in all measured life-history traits (Figure 3), suggesting that history may affect the evolutionary response to global warming. Higher thermal performance in southern *D. subobscura* populations has been shown by Porcelli et al. (2017). Longer-term experiments will evaluate if this incipient geographical differentiation in the thermal response expands further as populations evolve in the warming conditions.

### Cold stress effect is immediate and heat stress’ is transgenerational

In general, we found clear evidence for a plastic response to both cold and heat stress, despite general similarity of patterns between the different experimental populations. Exposure to colder temperatures led to an overall decline in fecundity of all populations. This effect was due, most likely, to lower metabolic rates, which may reduce oogenesis and lead to lower reproduction output, as previously observed (Simões et al. 2020). Unsurprisingly, life-long cold-exposed flies had better performance than flies that were only exposed to cold as adults, probably due to increased ovariole number during cold development (Moreteau et al. 1997) and showing evidence for beneficial cold acclimation in fecundity patterns (Huey et al. 1999; Simões et al. 2020).

Heat stress is known to significantly affect *D. subobscura* reproduction and, most importantly, the stage at which high temperature stress is experienced strongly influences reproductive performance (Porcelli et al. 2017; Simões et al. 2020; Santos et al. 2021). High developmental temperatures have shown to cause within-generation, negative carry-over effects in ectotherms (Klockmann et al. 2017; Porcelli et al. 2017; Cao et al. 2018; Iossa et al. 2019; Klepsatel et al. 2019), maybe due to the irreversible physiological and metabolic damage brought upon them (*e*.*g*., during gametogenesis). Conversely, adult exposure to warmer temperatures has, previously, shown to have little to no effect on fecundity and to increase the sexual maturity rate, leading to earlier reproduction (Santos et al. 2021). Our experimental data corroborates this dual effect in control and warming-selected flies: high lifelong temperature led to negative transgenerational effects in productivity and, also, in progeny viability, despite the high levels of fecundity. Furthermore, the longer the flies were exposed to the heat stress during their life cycle, the higher the damage in viability. Interestingly, this is not the case for fecundity, as flies with lifelong exposure to higher temperatures show higher egg production than those that experienced stress only in the development stage, suggesting some rescue in performance as a result of higher adult metabolic rates. The decoupling between fecundity and productivity/viability patterns further suggests that heat stress is (1) leading to the occurrence of male sterility; (2) affecting to a higher degree egg quality rather than egg production; 3) increasing juvenile mortality. In general, our results confirm the previously noted vulnerability of the insects’ development stage to climate change and its associated heat waves (Klockmann et al. 2017).

### Final considerations

Adaptation to a rapidly changing environment is of great importance for a population’s fitness and fate. Here we report no evidence for short-term evolution in the elevation or shape of the reaction norms of *D. subobscura* populations under dynamic thermal environments. As such, evolutionary change seems not to happen fast enough to counteract the detrimental effects of global warming, nor giving ectothermic animals tools to fight more extreme thermal conditions. Evolution, by itself, seems not be the ‘silver bullet’ needed to successfully mitigate rapid climate change (Diamond and Martin 2020). As evolutionary biologists, we tend to see the natural world through the lens of adaptation, as natural selection is exceptionally powerful in granting fitness advantages that shape population persistence and biological diversity. Nevertheless, it is becoming apparent that the recovery of populations (*i*.*e*., ‘evolutionary rescue’ (Bell 2017) may not occur quickly enough to keep up with the fast pace of current environmental change. A better understanding of the biology of endangered species will involve looking at their evolution with an open mind regarding these issues.

## Acknowledgments

The authors thank Inês Fragata for help in the graphic presentation of the data. This study is financed by Portuguese National Funds through ‘Fundação para a Ciência e a Tecnologia’ (FCT) within the projects PTDC/BIA-EVL/28298/2017 and cE3c Unit FCT funding project UIDB/BIA/00329/2020. PS and ASQ are funded by national funds (OE), through FCT, in the scope of the framework contract foreseen in the numbers 4, 5 and 6 of the article 23^rd^, of the Decree-Law 57/2016, of August 29, changed by Law 57/2017, of July 19. MS is funded by grants CGL2017-89160P from Ministerio de Economía y Competitividad (Spain; co-financed with the European Union FEDER funds), and 2017SGR 1379 from Generalitat de Catalunya.

## Authors Contributions

MAS participated in data collection and analysis, discussion of experimental data and wrote the first draft of the manuscript; A.C-S and ASQ participated in data collection, discussion of experimental data and manuscript writing; MS participated in discussion of experimental data and manuscript writing; MM and PS participated in data collection and analysis, discussion of experimental data and manuscript writing.

## Notes

### Competing Interest Statement

The authors have declared no competing interest.

