## Supplementary Material for "No evidence for short-term evolutionary response to a warming environment in Drosophila"

---

**Affiliations:** [a] cE3c – Centre for Ecology, Evolution and Environmental Changes, Faculdade de Ciências, Universidade de Lisboa, Lisboa, Portugal; [b] CESAM, Centre for Environmental and Marine Studies, Universidade de Aveiro and Faculdade de Ciências, Universidade de Lisboa, Lisboa, Portugal; [c] Departament de Genètica i de Microbiologia, Grup de Genòmica, Bioinformàtica i Biologia Evolutiva (GBBE), Universitat Autònoma de Barcelona, Spain. **Emails:** MAS * – corresponding author,; AC-S -; ASQ –; MS -; MM –; PS –

#### Supplementary Material

##### Supplementary Figures

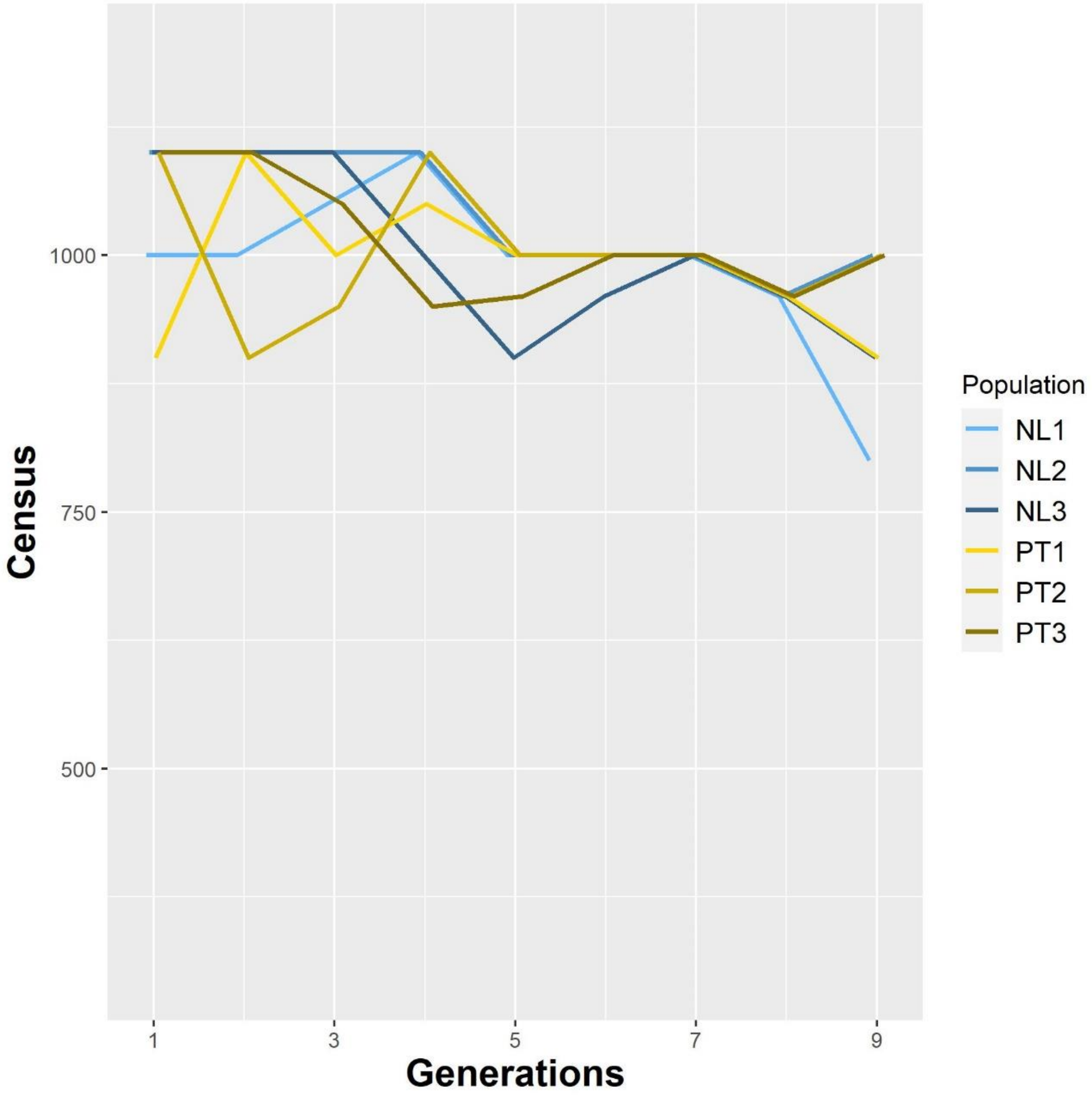


*Figure S1a – Census size of the control populations*

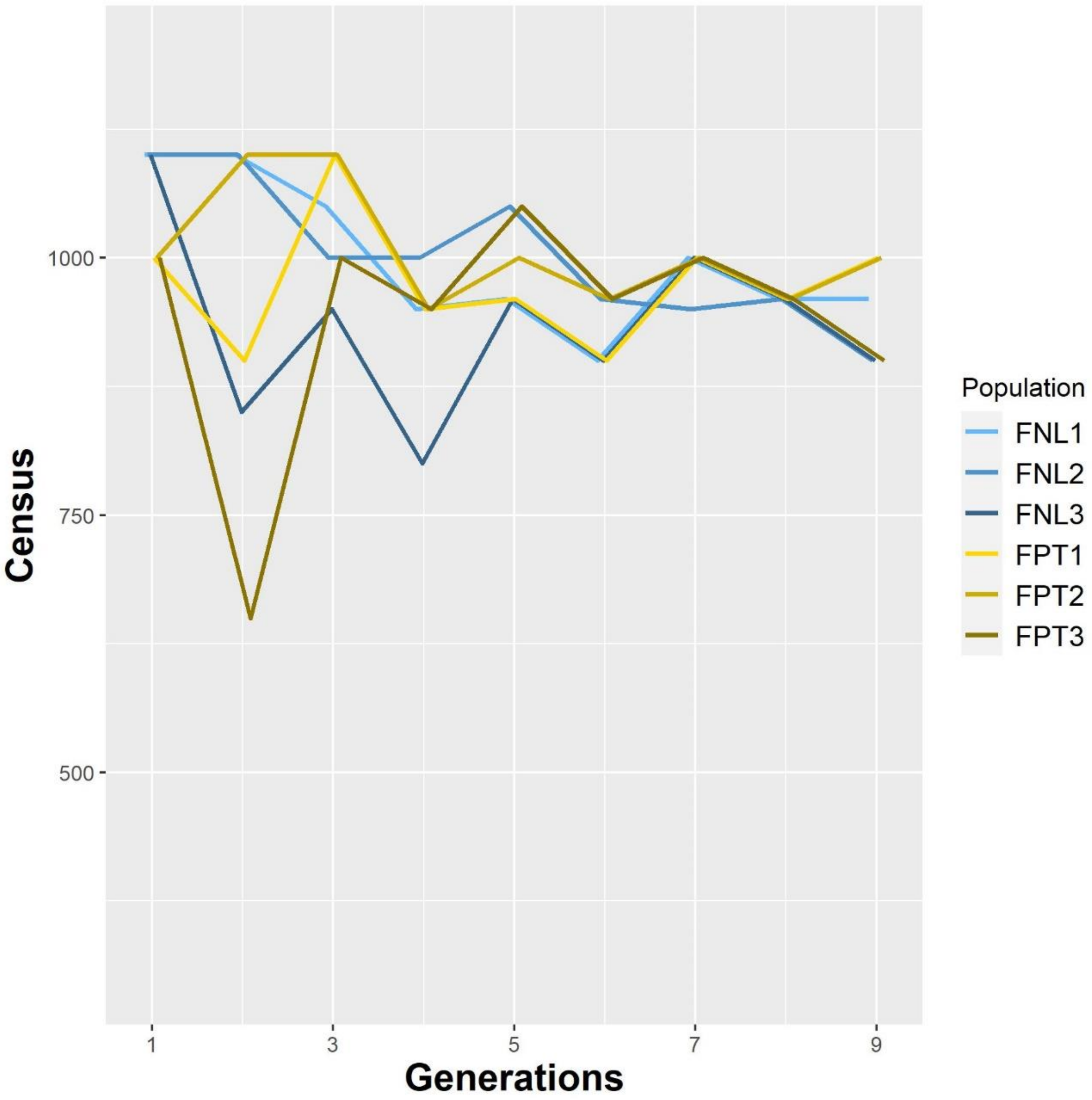


*Figure S1b – Census size of the fluctuating populations*

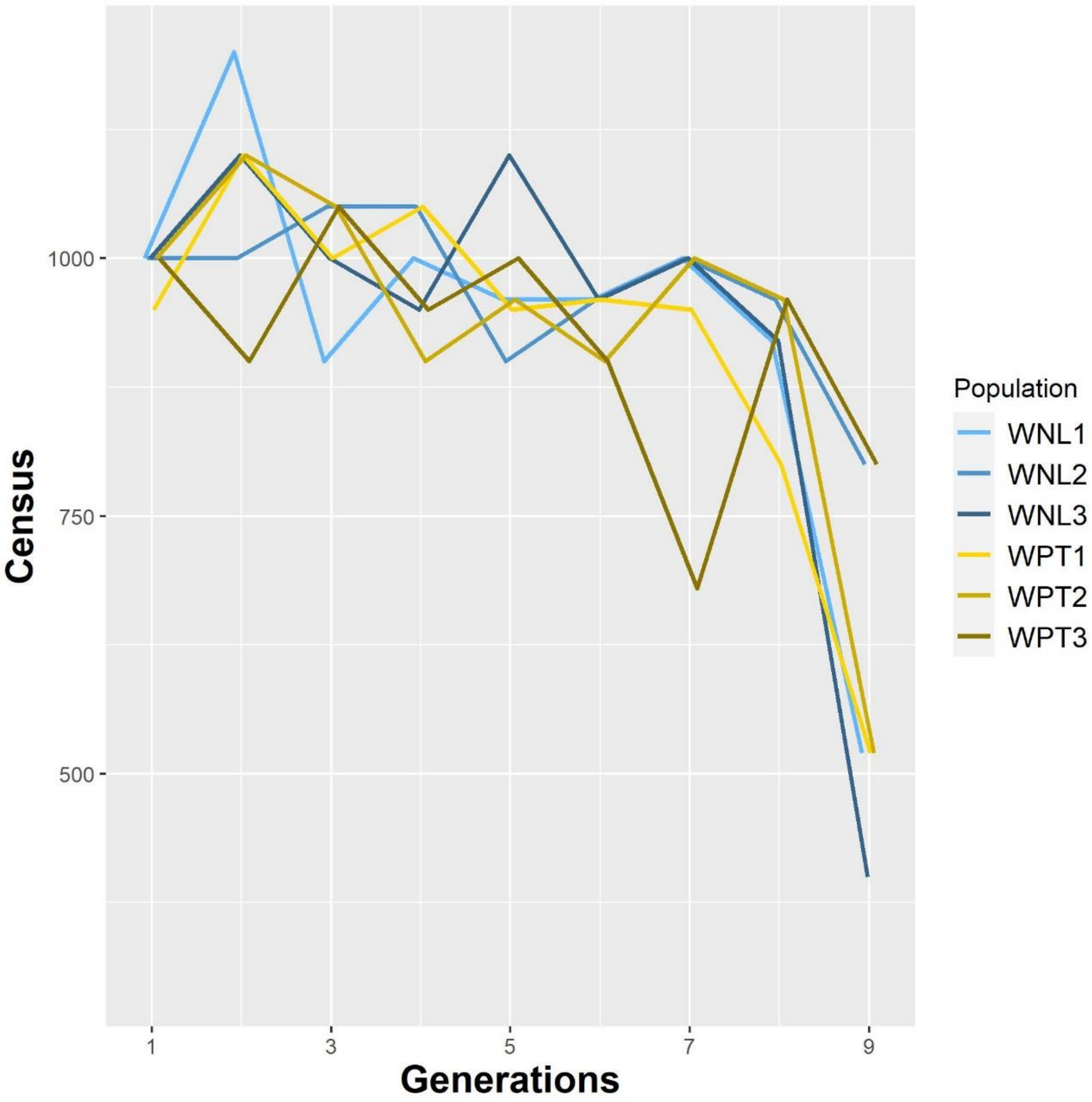


*Figure S1c – Census size of the warming populations*

*Figure S2*

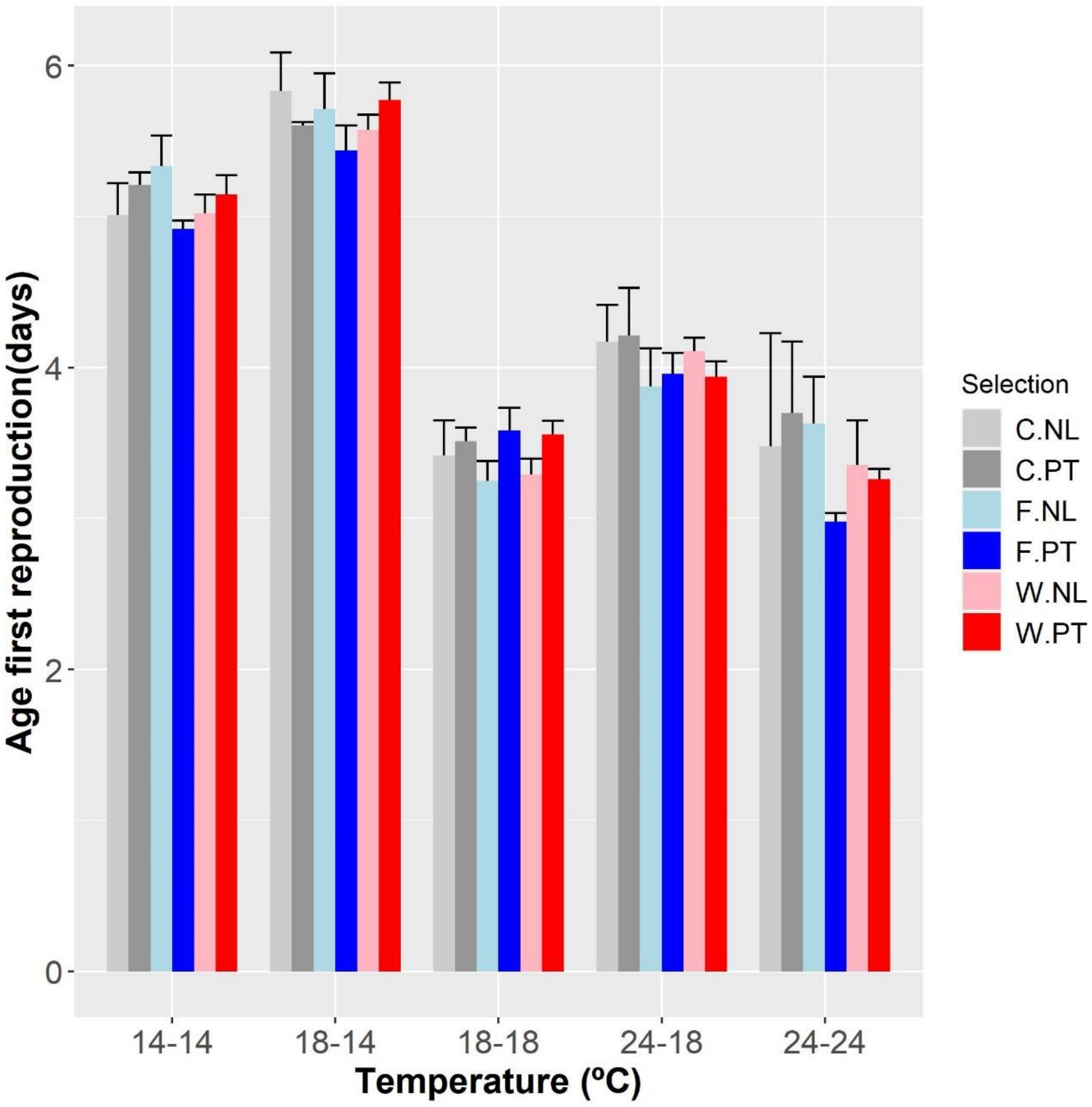

Age first reproduction(days)
0
2
4
6
14-14
18-14
18-18
24-18
24-24
Temperature (ºC)
Selection
C.NL
C.PT
F.NL
F.PT
W.NL
W.PT

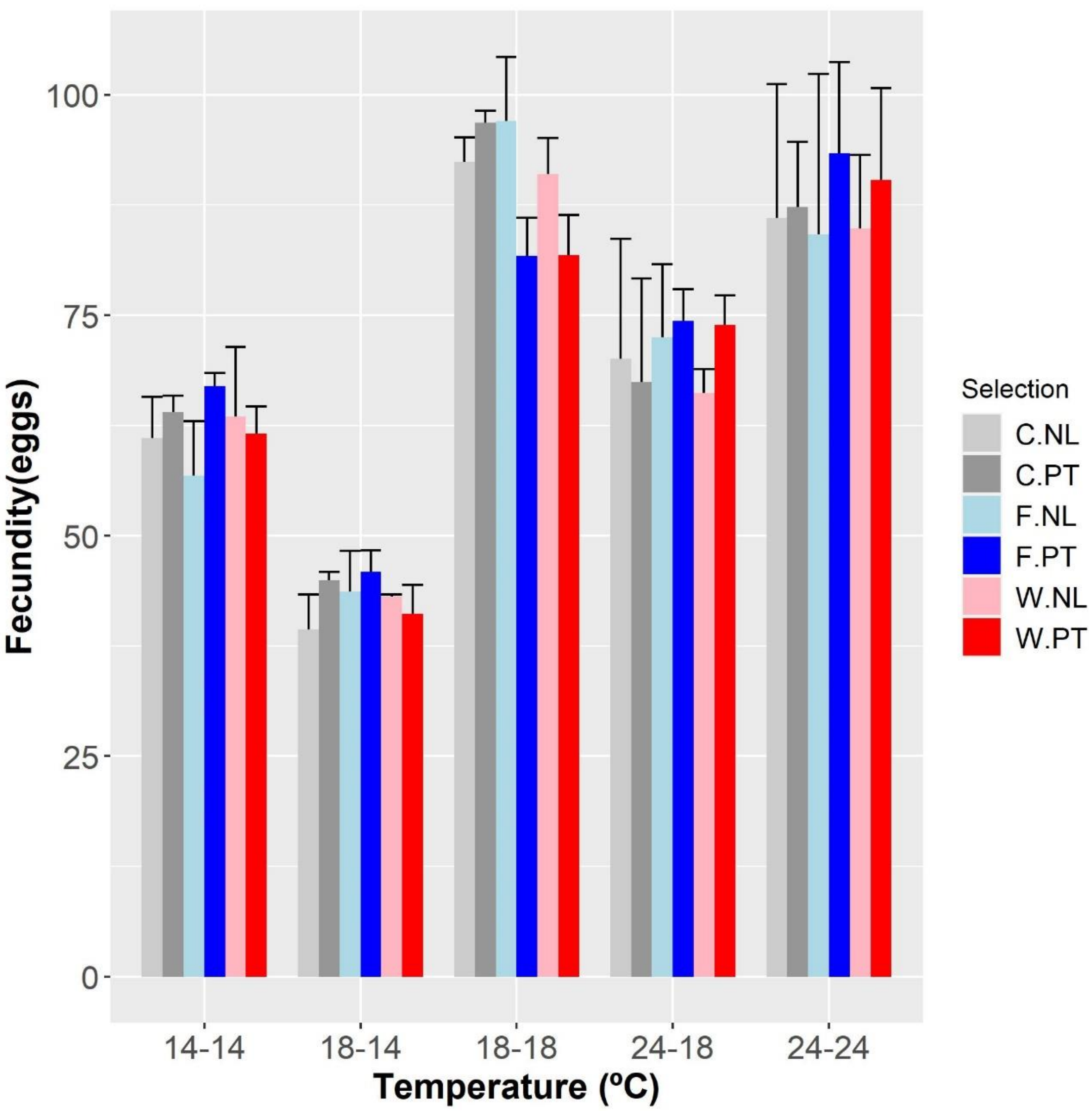
Fecundity(eggs)
100
75
50
25
0
14-14
18-14
18-18
24-18
24-24
Temperature (ºC)
Selection
C.NL
C.PT
F.NL
F.PT
W.NL
W.PT

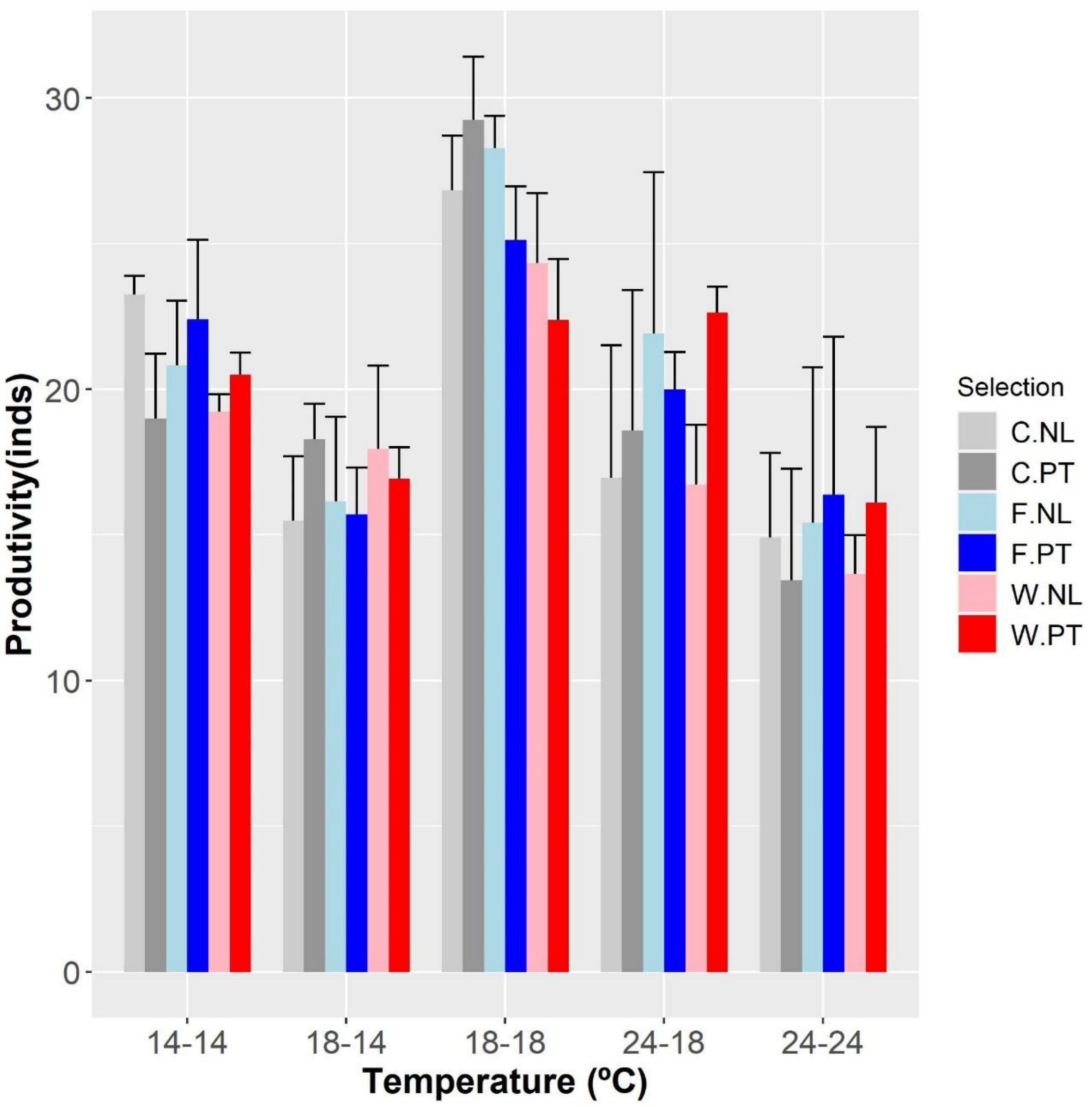
30
20
10
0
Produtivity(inds)
14-14
18-14
18-18
24-18
24-24
Temperature (ºC)
Selection
C.NL
C.PT
F.NL
F.PT
W.NL
W.PT

34

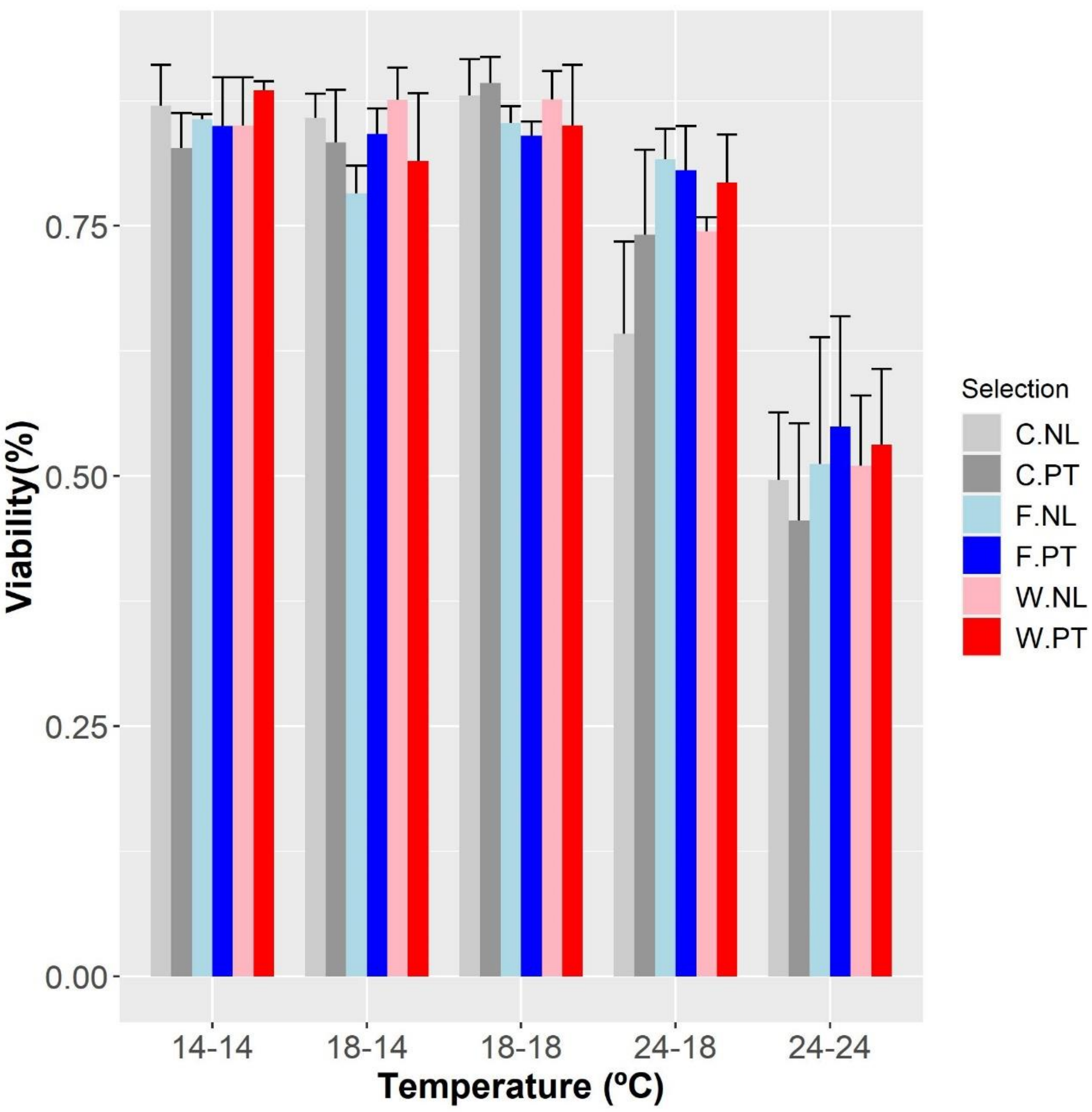


*Figure S2 – Thermal performance for populations of the different thermal regimes in all 5 temperature treatments. A) Age of first reproduction, B) Fecundity, C) Productivity, D) Juvenile Viability*

### Supplementary Tables

Table S1 - Differences in reaction norms between fluctuating and control populations. (14-14; 18-18; 24-24 treatments)

| Trait | Model parameters | $F_{(df1, df2)}$ |
|---|---|---|
| Age of First Reproduction (A1R) | History | $F_{1,4}$ = 0.015 n.s. |
| | Selection | $F_{1,4}$ = 0.549 n.s. |
| | Temp | $F_{2,8}$ = 49.469 *** |
| | History*Selection | $F_{1,4}$ = 0.654 n.s. |
| | History*Temp | $F_{2,8}$ = 0.653 n.s. |
| | Selection*Temp | $F_{2,8}$ = 0.846 n.s. |
| | History*Selection*Temp | $F_{2,8}$ = 2.857 n.s. |
| Fecundity | History | $F_{1,4}$ = 0.065 n.s. |
| | Selection | $F_{1,4}$ = 0.143 n.s. |
| | Temp | $F_{2,8}$ = 15.651 ** |
| | History*Selection | $F_{1,4}$ = 0.055 n.s. |
| | History*Temp | $F_{2,8}$ = 0.650 n.s. |
| | Selection*Temp | $F_{2,8}$ = 0.424 n.s. |
| | History*Selection*Temp | $F_{2,8}$ = 1.914 n.s. |
| Produtivity | History | $F_{1,22}$ = 0.147 n.s. |
| | Selection | $F_{2,22}$ = 0.030 n.s. |
| | Temp | $F_{2,22}$ = 17.949 *** |
| | History*Selection | $F_{2,22}$ = 0.073 n.s. |
| | History*Temp | $F_{2,22}$ = 0.042 n.s. |
| | Selection*Temp | $F_{2,22}$ = 0.280 n.s. |
| | History*Selection*Temp | $F_{2,22}$ = 1.019 n.s. |
| Viability | F8 | $F_{1,19}$ = 7.210 * |
| | History | $F_{1,4}$ = 0.004 n.s. |
| | Selection | $F_{1,19}$ = 0.332 n.s. |
| | Temp | $F_{2,21}$ = 54.556 *** |
| | History*Selection | $F_{1,19}$ = 0.927 n.s. |
| | History*Temp | $F_{2,19}$ = 0.248 n.s. |
| | Selection*Temp | $F_{2,19}$ = 0.810 n.s. |
| | History*Selection*Temp | $F_{2,20}$ = 0.252 n.s. |

Note: significance levels: p> 0.05 n.s.; 0.05>p>0.01*; 0.01>p>0.001**; p<0.001 ***

Table S2 - Differences in reaction norms between warming and control populations. (14-14; 18-18; 24-24 treatments)

| Trait | Model parameters | $F_{(df1, df2)}$ |
|---|---|---|
| Age of First Reproduction (A1R) | History | $F_{1,4}$ = 0.253 n.s. |
| | Selection | $F_{1,4}$ = 0.646 n.s. |
| | Temp | $F_{2,8}$ = 52.022 *** |
| | History*Selection | $F_{1,4}$ = 0.062 n.s. |
| | History*Temp | $F_{2,8}$ = 0.057 n.s. |
| | Selection*Temp | $F_{2,8}$ = 0.410 n.s. |
| | History*Selection*Temp | $F_{2,8}$ = 0.293 n.s. |
| Fecundity | History | $F_{1,4}$ = 0.008 n.s. |
| | Selection | $F_{1,4}$ = 0.949 n.s. |
| | Temp | $F_{2,8}$ = 12.407 ** |
| | History*Selection | $F_{1,4}$ = 0.908 n.s. |
| | History*Temp | $F_{2,8}$ = 0.111 n.s. |
| | Selection*Temp | $F_{2,8}$ = 1.382 n.s. |
| | History*Selection*Temp | $F_{2,8}$ = 1.094 n.s. |
| Produtivity | History | $F_{1,22}$ = 0.041 n.s. |
| | Selection | $F_{1,22}$ = 1.951 n.s. |
| | Temp | $F_{2,22}$ = 26.690 *** |
| | History*Selection | $F_{1,22}$ = 0.460 n.s. |
| | History*Temp | $F_{2,22}$ = 0.249 n.s. |
| | Selection*Temp | $F_{2,22}$ = 1.589 n.s. |
| | History*Selection*Temp | $F_{2,22}$ = 1.517 n.s. |
| Viability | F8 | $F_{1,22}$ = 0.257 n.s. |
| | History | $F_{1,4}$ = 0.021 n.s. |
| | Selection | $F_{1,19}$ = 0.450 n.s. |
| | Temp | $F_{2,20}$ = 41.772 *** |
| | History*Selection | $F_{1,19}$ = 0.360 n.s. |
| | History*Temp | $F_{2,19}$ = 0.053 n.s. |
| | Selection*Temp | $F_{2,19}$ = 0.375 n.s. |
| | History*Selection*Temp | $F_{2,19}$ = 0.109 n.s. |

Note: significance levels: p> 0.05 n.s.; 0.05>p>0.01*; 0.01>p>0.001**; p<0.001 ***

Table S3 - Differences in performance between warming and control selection regimes for combinations of colder (a) or warmer (b) thermal treatments

a)

| Trait | Model parameters | $F_{(df1, df2)}$ - Selection | $F_{(df1, df2)}$ - Selection x Temperature | $F_{(df1, df2)}$ - History | $F_{(df1, df2)}$ - History x Temperature | $F_{(df1, df2)}$ - History x Selection x Temperature |
|---|---|---|---|---|---|---|
| Age of First Reproduction (A1R) | 18-14 vs 18-18 | $F_{1,4}$ = 0.125 n.s. | $F_{1,4}$ = 0.002 n.s. | $F_{1,4}$ = 0.411 n.s. | $F_{1,4}$ = 1.555 n.s. | $F_{1,4}$ = 0.673 n.s. |
| | 14-14 vs 18-18 | $F_{1,4}$ = 0.122 n.s. | $F_{1,4}$ = 0.004 n.s. | $F_{1,4}$ = 1.912 n.s. | $F_{1,4}$ = 0.009 n.s. | $F_{1,4}$ = 0.455 n.s. |
| | 14-14 vs 18-14 | $F_{1,4}$ = 0.213 n.s. | $F_{1,4}$ = 0.013 n.s. | $F_{1,4}$ = 0.279 n.s. | $F_{1,4}$ = 0.737 n.s. | $F_{1,4}$ = 2.416 n.s. |
| Fecundity (F6-8) | 18-14 vs 18-18 | $F_{1,4}$ = 5.128 n.s. | $F_{1,4}$ = 5.041 n.s. | $F_{1,4}$ = 0.012 n.s. | $F_{1,4}$ = 0.748 n.s. | $F_{1,4}$ = 0.696 n.s. |
| | 14-14 vs 18-18 | $F_{1,4}$ = 2.832 n.s. | $F_{1,4}$ = 2.869 n.s. | $F_{1,4}$ = 0.070 n.s. | $F_{1,4}$ = 0.176 n.s. | $F_{1,4}$ = 0.817 n.s. |
| | 14-14 vs 18-14 | $F_{1,4}$ = 0.000 n.s. | $F_{1,4}$ = 0.0002 n.s. | $F_{1,4}$ = 0.081 n.s. | $F_{1,4}$ = 0.093 n.s. | $F_{1,4}$ = 0.100 n.s. |
| Productivity | 18-14 vs 18-18 | $F_{1,14}$ = 2.024 n.s. | $F_{1,14}$ = 3.215 n.s. | $F_{1,14}$ = 0.147 n.s. | $F_{1,14}$ = 0.048 n.s. | $F_{1,14}$ = 0.010 n.s. |
| | 14-14 vs 18-18 | $F_{1,14}$ = 5.722 m.s. | $F_{1,14}$ = 1.925 n.s. | $F_{1,14}$ = 0.254 n.s. | $F_{1,14}$ = 0.486 n.s. | $F_{1,14}$ = 4.011 n.s. |
| | 14-14 vs 18-14 | $F_{1,14}$ = 0.091 n.s. | $F_{1,14}$ = 0.582 n.s. | $F_{1,14}$ = 0.068 n.s. | $F_{1,14}$ = 1.023 n.s. | $F_{1,14}$ = 3.994 n.s. |

b)

| Trait | Model parameters | $F_{(df1, df2)}$ - Selection | $F_{(df1, df2)}$ - Selection x Temperature | $F_{(df1, df2)}$ - History | $F_{(df1, df2)}$ - History x Temperature | $F_{(df1, df2)}$ - History x Selection x Temperature |
|---|---|---|---|---|---|---|
| Age of First Reproduction (A1R) | 24-18 vs 18-18 | $F_{1,4} = 0.720$ n.s. | $F_{1,4} = 0.282$ n.s. | $F_{1,4} = 0.174$ n.s. | $F_{1,4} = 0.980$ n.s. | $F_{1,4} = 0.593$ n.s. |
| | 24-24 vs 18-18 | $F_{1,4} = 0.631$ n.s. | $F_{1,4} = 0.432$ n.s. | $F_{1,4} = 0.136$ n.s. | $F_{1,4} = 0.061$ n.s. | $F_{1,4} = 0.432$ n.s. |
| | 24-24 vs 24-18 | $F_{1,4} = 0.748$ n.s. | $F_{1,4} = 0.251$ n.s. | $F_{1,4} = 0.000$ n.s. | $F_{1,4} = 0.074$ n.s. | $F_{1,4} = 0.060$ n.s. |
| Fecundity (F6-8) | 24-18 vs 18-18 | $F_{1,4} = 0.596$ n.s. | $F_{1,4} = 1.129$ n.s. | $F_{1,4} = 0.0002$ n.s. | $F_{1,4} = 0.212$ n.s. | $F_{1,4} = 1.797$ n.s. |
| | 24-24 vs 18-18 | $F_{1,4} = 0.883$ n.s. | $F_{1,4} = 5.292$ n.s. | $F_{1,4} = 0.004$ n.s. | $F_{1,4} = 0.163$ n.s. | $F_{1,4} = 5.078$ n.s. |
| | 24-24 vs 24-18 | $F_{1,4} = 0.035$ n.s. | $F_{1,4} = 0.003$ n.s. | $F_{1,4} = 0.062$ n.s. | $F_{1,4} = 0.017$ n.s. | $F_{1,4} = 0.204$ n.s. |
| Productivity | 24-18 vs 18-18 | $F_{1,14} = 0.458$ n.s. | $F_{1,14} = 2.584$ n.s. | $F_{1,14} = 0.949$ n.s. | $F_{1,14} = 0.735$ n.s. | $F_{1,14} = 1.115$ n.s. |
| | 24-24 vs 18-18 | $F_{1,14} = 1.275$ n.s. | $F_{1,14} = 2.326$ n.s. | $F_{1,14} = 0.043$ n.s. | $F_{1,14} = 0.005$ n.s. | $F_{1,14} = 1.386$ n.s. |
| | 24-24 vs 24-18 | $F_{1,14} = 0.348$ n.s. | $F_{1,14} = 0.075$ n.s. | $F_{1,14} = 0.927$ n.s. | $F_{1,14} = 0.545$ n.s. | $F_{1,14} = 0.002$ n.s. |
| Viability | 24-18 vs 18-18 | $F_{1,11.70} = 0.650$ n.s. | $F_{1,11.14} = 1.032$ n.s. | $F_{1,4.98} = 0.150$ n.s. | $F_{1,11.04} = 1.277$ n.s. | $F_{1,11.28} = 0.216$ n.s. |
| | 24-24 vs 18-18 | $F_{1,11.16} = 0.567$ n.s. | $F_{1,11.01} = 0.610$ n.s. | $F_{1,3.93} = 0.063$ n.s. | $F_{1,11.10} = 0.249$ n.s. | $F_{1,11.01} = 0.104$ n.s. |
| | 24-24 vs 24-18 | $F_{1,11.36} = 2.422$ n.s. | $F_{1,11.07} = 0.028$ n.s. | $F_{1,3.62} = 0.413$ n.s. | $F_{1,11.49} = 0.174$ n.s. | $F_{1,11.13} = 0.607$ n.s. |

Note: significance levels: $p > 0.05$ n.s.
